# User-friendly Oblique Plane Microscopy on a fully functional commercially available microscope base

**DOI:** 10.1101/2024.01.09.574832

**Authors:** George Sirinakis, Edward S. Allgeyer, Dmitry Nashchekin, Daniel St Johnston

## Abstract

In this work we present an Oblique Plane Microscope designed to work seamlessly with a commercially available microscope base. To support all the functionality offered by the microscope base, where the position of the objective lens is not fixed, we adopted a two-mirror scanning geometry that can compensate for changes to the position of the objective lens during routine microscope operation. We showed that within the expected displacement range of the 100X, 1.35 NA objective lens away from its designed position, and for most practical applications, there is no significant effect on the resolving power, or the fidelity of the 3D data produced by the microscope. Compared to the more traditional scan-lens/galvo-mirror combination, the two-mirror scanning geometry offers higher light-efficiency and a more compact footprint, which could be beneficial to all OPM designs regardless of the use of a commercial base or not.

## 1. Introduction

Light-sheet fluorescence microscopy (LSFM) is a rapidly advancing set of techniques for biological imaging that offer impressive temporal resolution and low phototoxicity [1]. The underlying principle relies on selectively illuminating a thin slice of the sample around the focal plane of the objective that is used to collect the emitted fluorescence. To achieve this, most implementations of LSFM rely on a second, orthogonal, objective lens to focus the excitation light into a thin plane or sheet. However, this orthogonal geometry imposes strict mechanical constrains that limit the selection of available high-NA objectives. More importantly, having two orthogonal high NA lenses in close proximity excludes the use of typical sample mounting methods (e.g. slides, dishes, multi-well plates) and limits potential imaging applications.

The above shortcomings can be avoided by using a variant of LSFM, termed Oblique Plane Microscopy (OPM), which uses a single, high NA, objective to deliver the light-sheet illumination and collect the emitted fluorescence simultaneously [2]. In this approach, a light-sheet is introduced at an oblique angle, with respect to the primary objective, illuminating a thin slice of the sample that extends above and below the primary objective’s focal plane. This illumination scheme appears unconventional given that objective lenses are designed to capture aberration-free images only from their image plane, the in-focus plane of the sample, while imaging the other, out-of-focus planes suffers from spherical aberrations which become progressively worse with distance from the focal plane [3]. However, these aberrations can be cancelled out with remote focusing and the use of an appropriately matched second objective to create an aberration-free image of the sample at the focal space of the second objective [4,5]. It is then possible to obtain an image of the oblique plane without significant compromises in resolution using a glass-tipped tertiary objective which is tilted so that its focal plane aligns with the oblique plane illuminated by the light sheet [6–8]. Finally, by scanning the light-sheet and de-scanning the corresponding emission light with the help of a galvo mirror, 3D volumes can be collected at high speeds without mechanically moving the sample or the objective [9,10].

To date, most OPM systems have been implemented without the use of a commercial microscope base [11]. This approach allows the conjugation of the pupil plane of the primary objective with the scan mirror and the pupil plane of the secondary objective by arranging the required tube and scan lenses in a 4f configuration. This arrangement offers a straight-forward way to achieve optimal remote focusing performance and ensures that the angle of the light sheet is invariant during scanning and always aligned with the focal plane of the tertiary objective [5,12].

Commercial microscope bases provide convenient platforms that accommodate various imaging modalities (e.g. Transmitted Light, Epifluorescence, Differential Interference Contrast) and components (e.g. oculars, stage top heaters, incubators, etc) and enable a wide range of imaging experiments in a user-friendly way that is accessible to non-experts. Therefore, it would be highly beneficial to incorporate fully functional microscope bases in OPM designs, especially when navigating more complex samples, since the oblique view supported by the OPM modality may not be as intuitive as the more familiar cartesian one. However, integration of commercial microscope bases into OPM systems requires greater care as not only the position of the tube lens inside the microscope base deviates from the 4f configuration but sample focusing is typically achieved by moving the primary objective lens. Depending on the sample and/or mounting method, it is likely that the primary objective will move away from its design position as the user selects a focal plane, disturbing the conjugation with the scan mirror. Under these suboptimal conditions, scanning the mirror will result in the excitation beam pivoting around a point away from the back pupil plane of the primary objective. Hence, a step in the scan mirror will result in both a translation of the oblique light sheet in the sample volume and a change in its angle, which produces 3D volumes with non-uniform illumination and poor signal-to-noise ratio.

To address this challenge, we present an OPM configuration built around a commercial microscope base that allows full functionality of the base without disabling its focus capabilities. Specifically, we adopted a scanning configuration that employs a pair of galvo mirrors positioned close to the primary image plane of the microscope base [13]. By tuning the ratio of the angular displacement between the two mirrors, it is possible to change the axial position of the pivot point of the excitation beam and track the pupil plane of the objective as it moves during focusing. Therefore, regardless of the final primary objective position, the light-sheet will maintain its alignment with the focal plane of the tertiary objective during scanning to enable optimal 3D volumetric imaging. This tunability is not supported in the most traditional scan-lens/mirror configurations or other scanning geometries with only one movable mirror [7,14]. Furthermore, compared to the more traditional scan-lens/mirror combination, the two-mirror scanning geometry proposed here offers higher light-efficiency and a more compact footprint, which could be beneficial to all OPM designs regardless of whether they use a commercial base or not.

We took an empirical approach to quantify the remote focusing performance, as the primary objective is displaced away from its optimal position during routine microscope operation. We used fluorescent beads immobilized on a cover-glass and in agarose gel to demonstrate that for most imaging scenarios accommodated by a 1.35NA, 100X silicon oil objective lens, distortions on the remote focusing volume due to axial displacements of the objective lens are below 0.5%, but we also discuss strategies to eliminate these distortions for more demanding applications.

Lastly, we demonstrate the performance of the microscope by using Drosophila egg chambers as a test sample and following the dynamics of microtubules through whole follicle cells.

## 2. Experimental Methods

### 2.1 Optical Setup

Three telescopes are the key components of the OPM microscope as shown in Figure 1. The first telescope resides in the Olympus IX73 microscope base and is comprised of an Olympus 100X, 1.35 NA silicone oil objective (UPLSAPO100XS) (OBJ1) and a 180 mm tube lens (TL1). The second telescope is arranged back-to-back with the first telescope and forms an aberration free 3D image of the sample space in the remote volume superintended by the objective lens of this second telescope (OBJ 2), an Olympus 40X, 0.95 NA air objective (UPLXAPO40X). Key to the performance of the remote focusing system is that the overall magnification between the primary objective (OBJ 1) space and the remote focusing objective (OBJ 2) space should equal the ratio of the refractive indices of the immersion media in the primary and remote focusing objective spaces, respectively. Given that our choice for the primary and remote focusing objectives is an Olympus 100X Silicone oil (n = 1.406), and an Olympus 40X Air objective, respectively, the ideal focal length for the tube lens of the remote focusing objective (TL 2, Fig. 1) should be 320 mm. As no 320 mm focal length lenses are commercially available, we adopted a tube lens assembly that consists of two commercially available achromatic lenses (Thorlabs, ACT508-500-A-ML, and ACT508-750-A-ML) with an effective focal length of 321 mm as described in Millet-Sikking et al [6]. Finally, the third telescope is tilted at 28° with respect to the optical axis of the remote focusing objective (OBJ2) and employs a glass tipped objective lens (ASI, AMS-AGY, 54-10-5, v1.1) (OBJ3). This tertiary objective lens is mounted on a linear translation stage (Newport, 9066-X-M) driven by a motorized adjuster (Thorlabs, PIA13) to relay fluorescence signal from individual planes within the remote focusing volume through a quad-band emission filter (Chroma, ZET405/488/561/647m) to an sCMOS camera (Hamamatsu, Orca-Flash 4.0 v3) with the help of a 200 mm tube lens (Thorlabs, TTL200-A).

**Figure 1.**
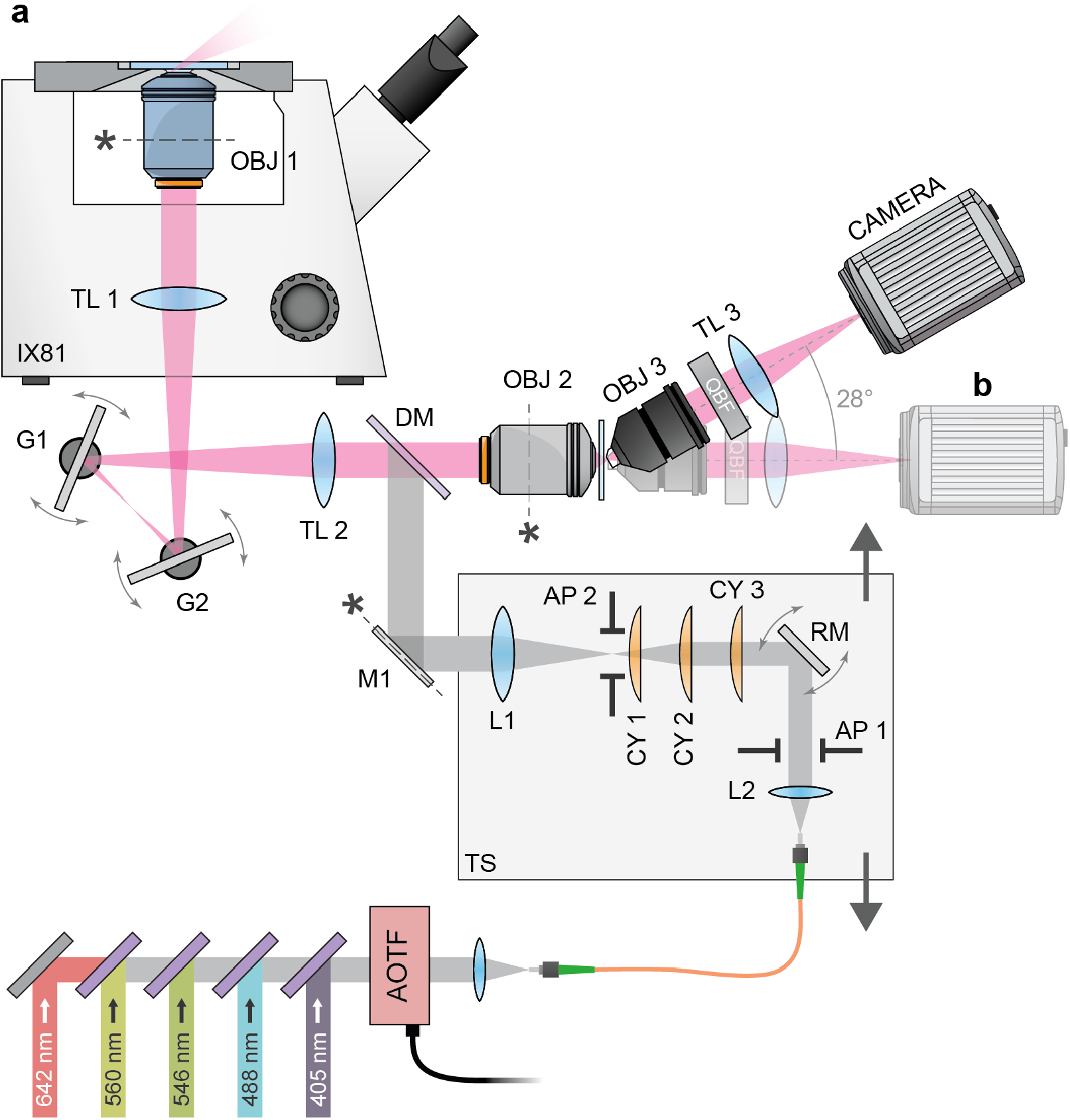
Experimental setup. **(a)** Excitation laser light is combined with a series of dichroic mirrors, coupled into a single mode fiber and collimated with L2, passes through adjustable slits (AP 1) for light sheet NA control and reflects off a 16 kHz resonance mirror (RM) for de-shadowing at the sample. The beam passes a set of 3 cylindrical lenses. CY 1 and CY 3 act as a telescope and adjust the beam size in one dimension while CY 2, rotated to be orthogonal with CY 1/3, creates a focused (light sheet) at AP 2. AP 2 controls the light sheet (field of view) extent at the sample. Next, light is focused (collimated) in one dimension with L1 before reflecting off a mirror (M1) conjugated to the primary objective (OBJ 1) back focal plane for controlling the light sheet position at the sample. Components L2 to L1 are all mounted on a translation stage (TS) that allows the excitation beam to be shifted, causing the position to move in OBJ 1’s back aperture and the light sheet to tilt. Excitation laser light is coupled into the common path with a quadband dichroic mirror (DM) and then focused by tube lens TL 2 before reflecting from two galvanometer mirrors (G2 and G1) and entering the side port of the microscope base (IX83). Finally, excitation light is focused in one dimension and collimated in the other by the microscope base’s tube lens TL 1 before entering the primary objective lens. Fluorescence is collected by the same primary objective (OBJ 1) and follows the same path back to dichroic mirror DM while being descanned by G1 and G2. After separating from excitation laser light, fluorescence enters the remote focusing objective lens (OBJ 2) and forms a focal volume at the front focal plane of OBJ 2. Finally, the tertiary objective (OBJ 3) is positioned with its front glass surface at the remote focusing plane matched to the light sheet at the primary objective tilted imaging plane. OBJ 3 is tiled at 28° and the subsequent quadband emission filter (OBF), tube lens (TL 3), and camera follow this tilted path. **(b)** The tertiary objective, tube lens, and camera are placed on a straight path for initial remote focusing characterization, setup and diagnostic purposes.

Light sheet excitation is provided by, five excitation lasers at 405 (Coherent, Obis 405 nm LX), 488 (Coherent, Obis 488 nm LS), 546 (MPB Communications, 2RU-VFL-P-1000-546-B1R), 560 (MPB Communications, 2RU-VFL-P-2000-560-B1R) and 642 nm (MPB Communications, 2RU-VFL-P-2000-642-B1R), all coupled into a single mode optical fibre (Thorlabs, PM-S405-XP Custom). Laser light emitted from the fibre is collimated with an f = 30 mm achromat (Thorlabs, AC254-030-A) and passes through an adjustable slit (Thorlabs, VA100CP) to control the beam size in one dimension and, subsequently, the light sheet numerical aperture (NA). Next, collimated light reflects from the surface of a 16 kHz resonance mirror (EOPC, SC-30). The mirror surface is imaged onto the sample plane by downstream optics for the purpose of pivoting the light sheet, in plane, to de-shadow during imaging [15]. After reflecting from the 16 kHz resonance mirror, the excitation beam passes through three cylindrical lenses. The first (f = 25 mm, Edmund Optics, 69-717) and last (f = 100 mm, Thorlabs, LJ1567RM-A) cylindrical lenses act on one dimension of the excitation beam as a telescope and expand the beam diameter, only in that dimension, by a factor of 4 and setting the width of the light sheet upon exiting the primary objective lens at the sample. The middle cylindrical lens (f=150mm, Thorlabs, LJ1629RM-A) is oriented orthogonally to the first and third and is responsible for creating the light sheet. After the set of cylindrical lenses, an adjustable iris further controls the extent of the light sheet (width) at the sample plane to match the desired field of view (FOV). These elements are all mounted on a translating platform whose position is adjusted to displace the excitation light in the primary objective back aperture and set the final light sheet angle at the sample. The excitation beam reflects from a mirror that is conjugated to the primary objective back focal plane. This mirror allows the position of the light sheet to be adjusted, at the sample plane, without changing the angle. Next, the excitation beam reflects from a quad-band dichroic mirror (Chroma, ZT405/488/561/640rpcv2), passes through the 321 mm tube lens assembly (Fig. 1, TL 2), and reaches two galvanometer (galvo) mirrors (Thorlabs, QS30X-AG). Both galvo mirrors are connected to separate analogue outputs from an FPGA based data acquisition (DAQ) board (National Instruments, PCIe-7852R) and are actuated synchronously to scan the excitation light sheet across the sample and de-scan the corresponding fluorescence signal. Microscope control and data collection are carried out within the LabView environment using custom control software.

### 2.2 Remote Focusing Volume Characterization

To evaluate the remote focusing performance, we used 100 nm green/yellow beads (Thermo Fisher, F8803) immobilized with Poly-L-lysine (Sigma, P4707-50mL) on a No 1.5 cover glass bottomed 8 well dish (ibidi, 80807) and immersed in silicone oil. To achieve uniform illumination of the beads across the FOV we used the epifluorescence module of the Olympus microscope base which was equipped with an Olympus filter cube (Olympus, U-MWIBA3) which selected the spectrum for green excitation of a mercury arc lamp (X-Cite, EXFO 120).

We mounted the remote focusing objective (OBJ 2) on a linear stage (Thorlabs, XR25P/M) that allowed displacement along its optical axis ± 5 mm away from its design position. This enabled us to systematically characterize how the performance of the remote focusing system is affected by axial misalignments between the primary and the remote focusing objective and experimentally confirm the optimal axial position of the latter.

We also replaced the tilted third telescope downstream of the remote focusing objective with one comprised of an Olympus 60X, 0.9 NA air objective (UPLFLN) and a 150 mm tube lens (Thorlabs, AC254-150-A-ML) set on a straight optical path (Fig. 1b). This configuration simplifies data analysis as the mapping between the primary and remote focusing volumes can be readily described in cartesian coordinates without any transformations.

Finally, we mounted the Olympus 60X air objective lens on a long travel range piezo walk stage capable of 14 mm travel with nanometer step sizes (Physik Instrumente, N-664.3A). In this way, we could probe different parts of the remote volume with high resolution regardless of the axial position of the remote focusing objective and collect bead Z-stacks for PSF measurements.

#### 2.2.1 PSF measurements

To measure the PSF we mounted the bead sample on the microscope’s XYZ stage (Applied Scientific Instrumentation, MS-2000 XYZ) with the Z piezo set at the centre of its range of motion. Then, the beads were epi-illuminated and brought into focus using the eye pieces and adjusting the focus knob (objective Z-position) of the microscope base. We consider this the design or reference position of the primary objective and we kept the primary objective at this position for all PSF measurements. Subsequently, we used the Z piezo of the microscope’s stage to position the beads at 8 different depths away from the focal plane of the primary objective at Z = -40, -30, -20, -10, 10, 20, 30, 40 μm. For each Z position of the bead sample in the primary objective space, a bead image was formed at a corresponding depth in the remote volume. We could then probe these parts of the remote volume with the help of the third telescope and collect a bead image stack by scanning the tertiary objective with 54 steps and a 150 nm step size. To measure the axial and lateral FWHMs of the PSF we used custom Matlab scripts. First, the frame with the best overall focus was determined based on the second derivate method presented by Pech-Pacheco *et al*., candidate bead locations were identified using a wavelet transform algorithm as described in Y. Zhang et al. and fit with a 2D Gaussian to determine the XY position for each bead across the FOV [16,17]. Subsequently, the Z profile for each bead at its fitted XY position was extracted from all the images in the stack and fit with a 1D Gaussian to determine the axial (Z) FWHM of the PSF and to identify the best focal plane for each bead. Finally, a second 2D Gaussian fit was performed at the best focal plane of each bead to determine the lateral (X, Y) PSF FWHM.

#### 2.2.2 Remote volume distortion due to axial misalignment

To investigate how the axial misalignment between the primary and remote focusing objective affects the PSF and the remote focusing volume, we kept the primary objective lens fixed at its design position and translated the remote focusing objective lens in 1 mm steps to cover a total range of ± 5 mm around its design position, which was determined from a Zemax model of the microscope. At each new axial position of the remote focusing objective lens, we performed the PSF analysis routine described in Section 2.2.1 to determine the PSF lateral and axial FWHM and the position of each bead, at different depths of the remote focusing volume.

We observed that as the axial misalignment between the primary and remote focusing objective lenses was increased, bead positions appeared to radially shift further from the image centre with increasing remote focusing depth. Furthermore, the amount of radial shift was observed to be proportional to the distance from the image centre.

To quantify this radial shift at each remote focusing depth, we converted the position of each bead into polar coordinates (r, θ) and found the corresponding bead with the same θ value and smallest r separation in the centre of the remote focusing volume, where the bead sample is positioned at the focal plane of the primary objective (Z=0). Subsequently, the radial bead displacement of all beads in the FOV was plotted as a function of the distance from the image centre and fit with a linear function to determine the rate of radial displacement at that depth of the remote focusing volume.

### 2.3 Scan mirror voltage calibration

To determine the optimal voltage ratio between the scanning mirrors as a function of the primary objective Z position, we used 100 nm diameter beads (Thermo Fisher, F8803) embedded in a 2% agarose gel. The primary objective was initially set to the lowest z position (0 mm) allowed by the microscope base and the 3D bead sample was brought into focusing using a custom Z stage insert that allowed the sample to be translated over a 25 mm Z range using a micrometre driven linear stage (Thorlabs, XR25C/M).

We used a light-sheet with NA = 0.18 to illuminate an oblique FOV with dimensions of 98 μm x 16 μm, set the voltage ratio between the scanning mirrors to a starting guess and performed 100 steps to scan a total length of 140 μm. For each step of the scanning mirrors, we collected an image and obtained the summed intensity value. Next, we normalized this intensity value to the average intensity of all the 100 images collected at each step of the mirrors, plotted the normalized intensity values as a function of step number and fit with a linear function to obtain the slope value for this voltage ratio.

Subsequently, we changed the voltage ratio between the scanning mirrors and imaged the same volume again to obtain the slope value for the new voltage ratio. In total we investigated 15 different voltage ratios from 1.013 to 1.041 and by plotting the corresponding slope values and performing a linear fit we predicted the voltage ratio that resulted in a slope of zero for the lowest primary objective position.

The primary objective lens was then moved up by +1 mm and the above procedure was repeated to obtain the voltage ratio that produces a zero slope for the new objective position. We investigated a total of 10 primary objective lens Z positions, with a step size of 1 mm, spanning the entire range of motion offered by the microscope base.

### 2.4 Microscope characterization oblique path

To measure the PSF in the oblique path we mounted the bead sample on the microscope’s XYZ stage with the Z piezo set at the centre of its range of motion. The beads were epi-illuminated and brought into focus using the eye pieces and adjusting the focus knob of the microscope base to bring the primary objective to its design position. Subsequently we switched to the OPM modality, used a 488 nm light-sheet with an NA of ∼0.1, and collected a volume of 77 × 90 x 26 μm by scanning the mirrors with a 117 nm step size along the X direction. Subsequently, we used the microscope stage’s Z piezo to position the beads at 4 different depths away from the focal plane of the primary objective at Z = -10, -5, 5, 10 μm. For each Z position of the bead sample, we imaged the same volume under identical imaging conditions. To analyse the beads and obtain the lateral and axial FWHMs we first deskewed the raw volumes to convert to cartesian coordinates and then followed the same analysis steps described in section 2.2.1. To obtain a summary image we merged the deskewed datasets into a single volume.

### 2.5 *Drosophila* Sample Preparation

*Drosophila* egg chambers were prepared as previously described with minor modifications [18]. Briefly, flies expressing the microtubule-associated protein, Jupiter, endogenously tagged with eGFP were fattened on dry yeast at 25°C for one day before dissection [19]. Their ovaries were dissected at room temperature in a drop of Voltalef oil 10S (VWR, 24627.188) on a cover glass (22x50, thickness No. 1.5, VWR, 631-0138) and individual egg chambers were isolated and positioned at the centre of the coverslip.

## 3. Results and Discussion

### 3.1 Remote focusing volume PSF characterization

To characterize the performance of the remote focusing system we worked on the straight optical path **(Fig. 1b)**. In this configuration both the primary and remote focusing objective volumes can be intuitively mapped in cartesian coordinates without any postprocessing. We began by positioning the fluorescent microspheres at the focal plane of the primary objective and measuring their corresponding lateral and axial FWHM across the available field-of-view (FOV). As expected, beads closer to the centre of the FOV exhibited slightly improved FWHMs (255 ± 10 nm laterally and 688 ± 38 nm axially) compared to the edges (272 ± 14 nm laterally and 764 ± 49 nm axially) (Fig. 2a). But, on average, throughout a circular area of 200 μm in diameter we obtained a relative uniform performance with FWHMs of 264 ± 15 and 730 ± 49 nm (mean ± standard deviation, n = 974) for the lateral and axial dimensions respectively.

**Figure 2.**
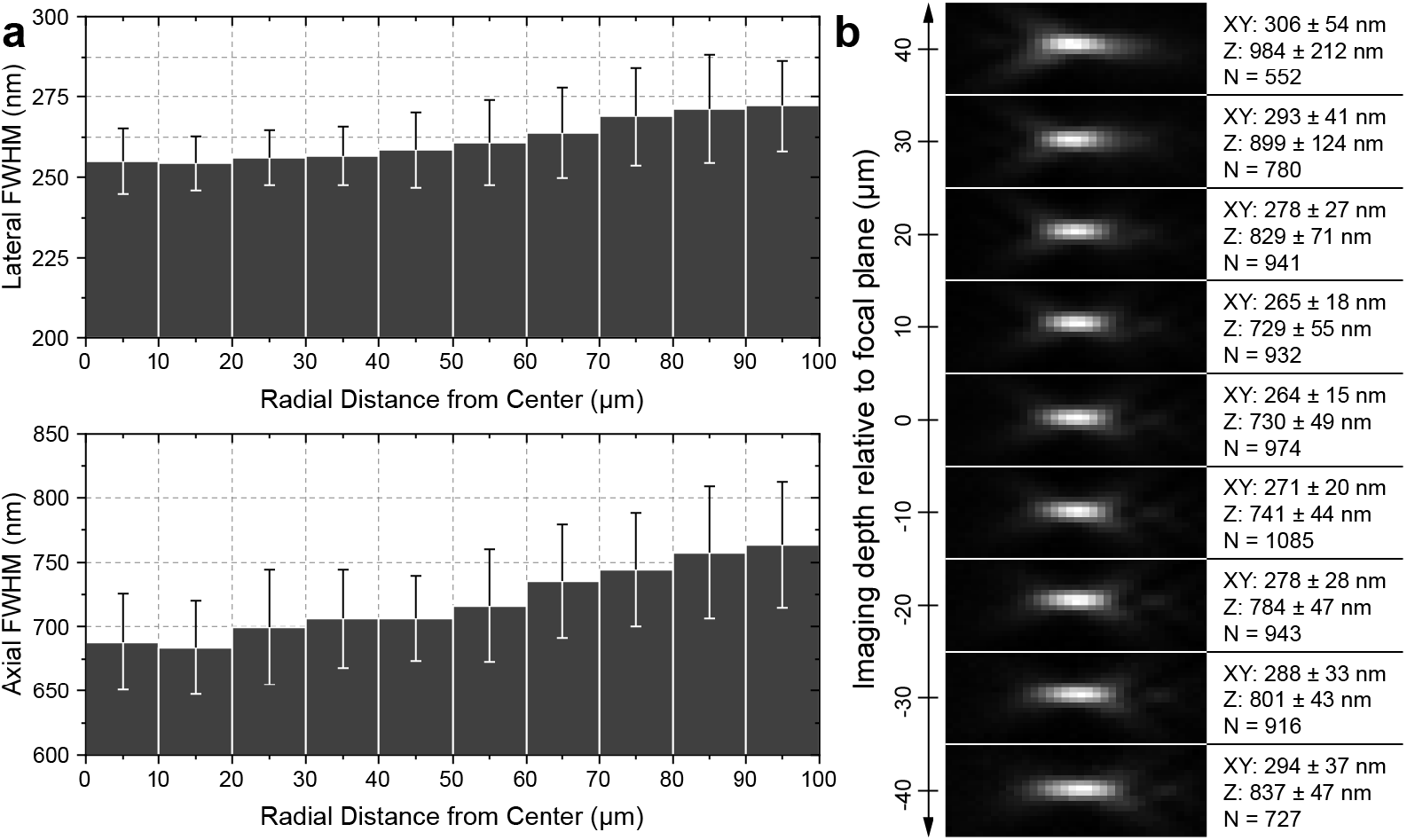
Remote focusing PSF characterization. (**a**) Lateral (XY) and axial (Z) PSF FWHMs as a function of radial distance from the image centre with a bead sample at the primary objective focal plane. (**b**) Representative PSF cross sections and mean fitted XY, Z FWHM and number of beads with the bead sample at nine Z positions. Each example PSF’s grey scale is set to its min/max.

To provide some context for the above measurements, we imaged the same beads with the epifluorescence modality of the Olympus microscope base, via the camera port on the eyepiece adaptor, and obtained 238 ± 3 nm and 593 ± 30 nm (mean ± standard deviation, n = 795) for the lateral and axial FWHMs respectively. Given that in the epifluorescence configuration, only the primary objective and its tube lens inside the microscope base affect the size and shape of the PSF, we reasoned that the epifluorescence values for the axial and lateral FWHMs would provide a good experimental reference for the best performance that can be expected from the remote focusing system. Therefore, the good agreement between the axial and lateral FWHMs measured with the remote focusing system and the epifluorescence module indicate that no significant aberrations were introduced by the additional optical components of the remote focusing system, which exhibits strong imaging performance.

To study how the PSF changes throughout the remote focusing volume, we kept the primary objective fixed at its designed position, while the bead sample z-position was discretely varied from -40 to +40 μm in 10 μm steps. Bead stacks were collected by scanning the tertiary objective as described in section 2.2.1. Representative PSF cross sections are shown in **Fig. 2b**. The PSF of the system retained its overall shape over the investigated depth range indicating that no significant aberrations were introduced by the remote focusing system. However, we noticed a small increase in the lateral size and an elongation along the axial direction of the PSF which become more pronounced with axial distance from the centre of the remote focusing volume. These changes on the size of the PSF indicate the presence of residual spherical aberrations that are not eliminated by the remote focusing system, possibly due to deviations of the magnification between the object space and the remote focusing volume from its optimal value [20]. Nonetheless, the shape of the PSF does not change meaningfully for practical applications, within the investigated range, and delivers slightly better performance over volumes extending ± 20 μm away from the focal plane of the primary objective.

A key incentive for using a microscope base is the ability to navigate the sample and focus on the plane of interest. This is typically achieved by moving the primary objective axially. Although this type of motion does not affect standard microscope operation it may disturb the axial alignment between the primary and remote focusing objectives. To investigate how axial misalignment between the primary and remote focusing objectives would impact the PSF, we fixed the primary objective at its nominal position and mounted the remote focusing objective on a translation stage that supported a large axial displacement range of ± 5 mm as described in section 2.2.2. Given that the primary objective has a working distance of 200 μm and this system is built around an IX81 base with a microscope stage equipped with inserts for slides and dishes, the deviations from the nominal primary objective position during routine operation are not expected to exceed ±1 mm and displacing the remote focusing lens by ± 5 mm already covers this range (based on the axial magnification). Average X, Y and Z FWHM values for each sample z-position and remote focusing objective position are presented in **Fig. 3a**. Despite axially misaligning the remote focusing objective by ± 5 mm, the PSF maintained its overall shape, and only for axial misalignments larger than ± 2.5 mm and depths larger than 30 μm away from the focal plane, we observed a more pronounced increase in the lateral size of the PSF and an elongation along its axial dimension **Fig. 3b**. Although, these changes in the PSF size indicate the presence of more pronounced spherical aberrations and consequently a reduction in the volume over which the remote focusing system exhibits good performance, the PSF does not seem to be very sensitive to axial misalignments between the primary and remote focusing objectives. In fact, the PSF retained its overall shape and size over a volume extending ± 20 μm away from the focal plane of the primary objective, regardless of the axial misalignment between the primary and remote focusing objectives within the investigated range.

**Figure 3.**
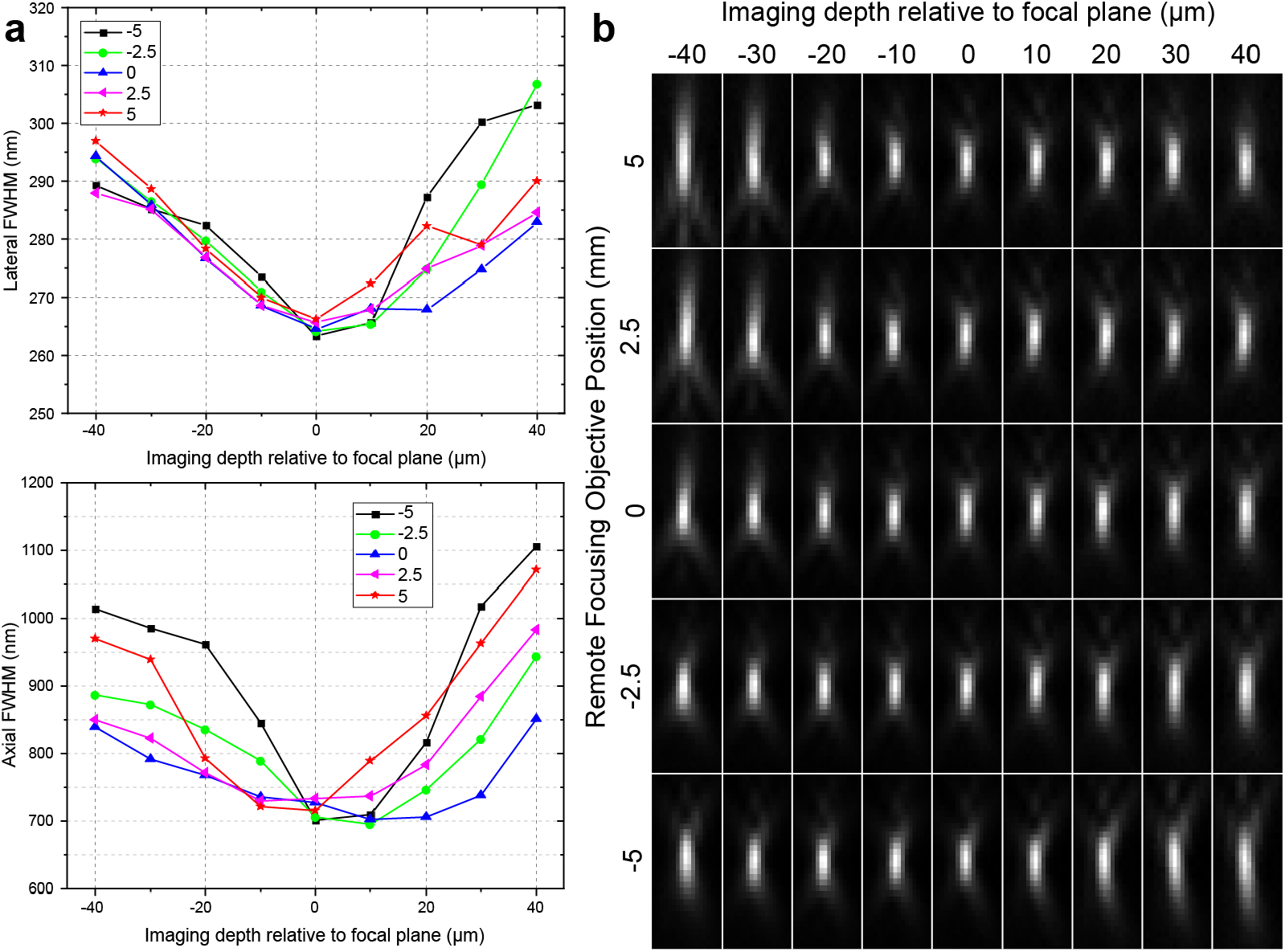
Remote focusing PSF characterization for different levels of axial misalignment between the primary and remote focusing objectives. (**a**) Lateral and axial FWHM of the PSF as a function of imaging depth for different axial positions of the remote focusing objective relative to its design position. (**b**) The corresponding example PSF XZ cross sections.

### 3.2 Remote focusing volume distortion characterization

Another effect of the axial misalignment between the primary and remote focusing objectives is a distortion of the remote focusing volume, which is perceived as a change in magnification with sample Z-depth [21]. To investigate the amount of this distortion in our system, we used the same sample and experimental procedure, as above for the PSF measurements, but in this case for each axial position of the remote focusing objective, we first identified all the beads in an image recorded with the sample positioned at the Z = 0 focal plane of the primary objective, and subsequently we tracked the beads through the rest of the images that were recorded above and below the focal plane of the primary objective as described in section 2.2.2. We then measured the bead’s lateral displacements relative to their position at the focal plane and calculated the rate of radial separation, which provides a good description of the changes in magnification introduced by the remote focusing system at different depths of the remote focusing volume and for different degrees of axial misalignment between the primary and remote focusing objective. The results are summarized in **Fig. 4a**. The magnification remains relatively constant throughout the remote focusing volume for axial positions of the remote focusing objective near its design position. As the remote focusing objective moves to axial positions further above its design position we observe increasing levels of distortion in the remote focusing volume with the apparent magnification increasing for planes above and decreasing for planes below the focal plane and vice versa for axial positions below the design position. The design position of the remote focusing objective was estimated from a Zemax model of the microscope, but without access to an optical model for the Olympus objective and tube lens and a CAD model for the IX81 base. Given these limitations, we considered the design position of the remote focusing objective as a good initial guess and proceeded to determine its optimal axial position experimentally. By plotting the rate of the magnification change versus axial position and performing a linear fit we obtained the optimal axial position of the remote focusing objective lens at ∼0.4 mm away from its initial design position (**Fig. 4b**).

**Figure 4.**
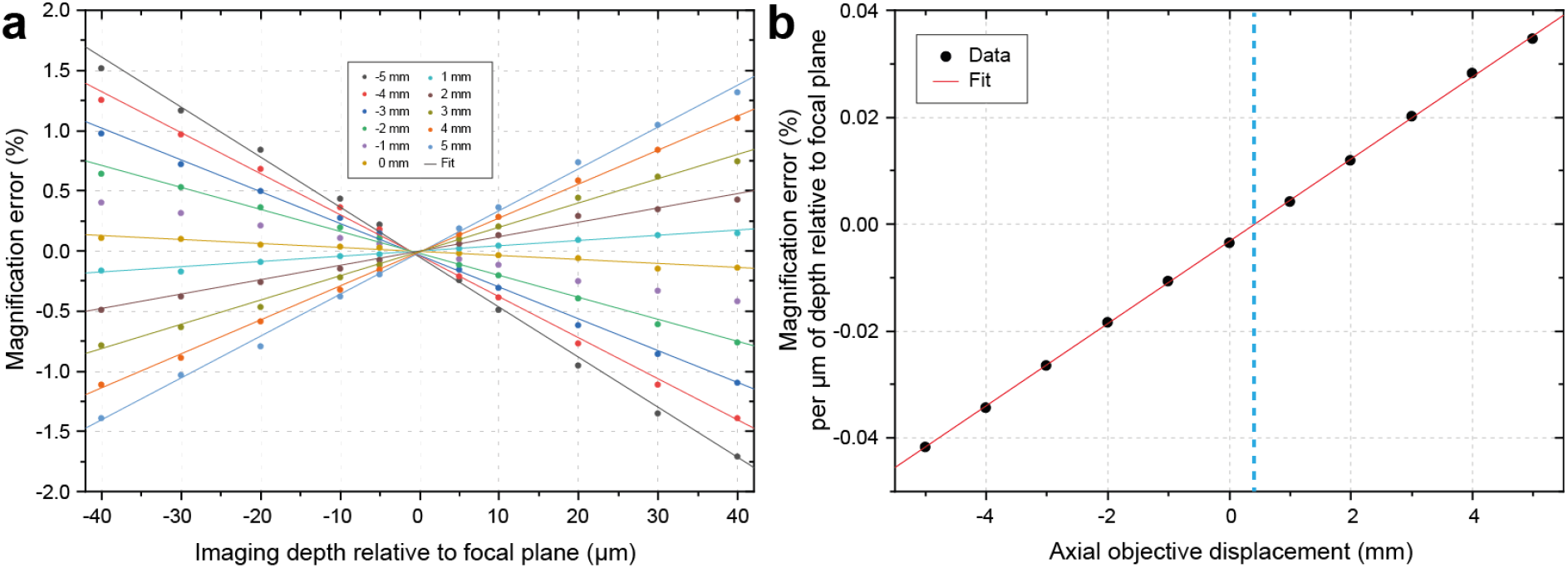
Remote focusing volume distortion characterization. **(a)** The percent error in magnification is plotted as a function of imaging depth relative to focal plane for different axial positions of the remote focusing objective. **(b)** The magnification error rate is plotted as a function of the axial objective displacement to obtain the optimal position of the remote focusing objective where the magnification remains constant throughout the remote volume.

For our applications, where we will operate the OPM microscope at a tilt angle of ∼30° and employ light sheets with lengths up to ∼40 μm, the working depth of the primary objective space is not expected to exceed ± 10 μm [7,22]. Furthermore, given that the primary and remote focusing objectives are conjugated via their corresponding tube lenses with focal lengths of 180 and 321 mm respectively, the amount of misalignment introduced during routine microscope operation, where the primary objective could be displaced by up to ± 1 mm from its designed position while the remote focusing objective remains fixed, is equivalent to the scenario where the primary objective is fixed and the remote focusing objective is displaced by ± 1.8 mm. Consequently, under routine microscope operation, the PSF of the system will not be significantly affected and distortions of the remote focusing volume are expected to remain at levels below 0.5% (**Fig. 4a**). For most practical applications, this distortion is negligible and does not warrant additional modifications to the optical path to compensate for it. However, for other scenarios that require longer travels of the primary objective and/or employ light sheets that illuminate volumes with greater depths, the axial misalignment between the primary and remote focusing objectives may compromise the performance of the microscope and the fidelity of the volumetric data. One way to address this issue, that also allows for uncompromised operation of the microscope base, would be to insert a pair of glass wedges in the optical path between the remote focusing objective and its tube lens (**Supplementary Fig.1**). By changing the thickness of the glass in the optical path it is possible to compensate for the misalignment introduced by the motion of the primary objective during microscope operation and ensure that the primary and remote focusing objectives always remain conjugated so that optimal performance is maintained, regardless of the position of the primary objective. A detailed description of the proposed optical path along with drawings for the wedges are provided in the appendix.

### 3.3 Microscope characterization, oblique path

Next, we moved to the oblique path by inserting downstream of the remote focusing objective a third telescope comprised of the glass-tipped objective and a 200 mm tube lens (**Fig. 1**). The optimum choice of illumination angle and, consequently, the tilt of this third telescope with respect to the focal plane of the remote focusing objective has been discussed in detail by Millet-Sikking et al., where it was shown that for an 1.35 NA silicone oil objective a ∼30° tilt offers a good balance between optical sectioning and size of illuminated volume [6,7]. In our system the tilt angle was 28 degrees.

To characterize the performance of the overall system we first imaged 100 nm yellow beads as described in section 2.4 (**Fig. 5a**). Near the focal plane of the primary objective, we measured average PSF FWHMs of ∼290 nm and ∼730 nm in the lateral and axial dimension respectively, in good agreement with our measurements on the straight path (**Fig. 5b**). But, as the distance from the focal plane increased these values also increased to ∼315 and ∼950 nm for the lateral and axial FWHM respectively at depths of ± 10 μm. Depths further away from the focal plane of the primary objective will be imaged further away from the centre of the tertiary objective and are expected to experience more aberrations. This is consistent with our observations and could explain the increase in the axial and lateral FWHMs. Despite these changes, the PSF maintains its overall shape and size and delivers strong performance within the investigated range.

**Figure 5.**
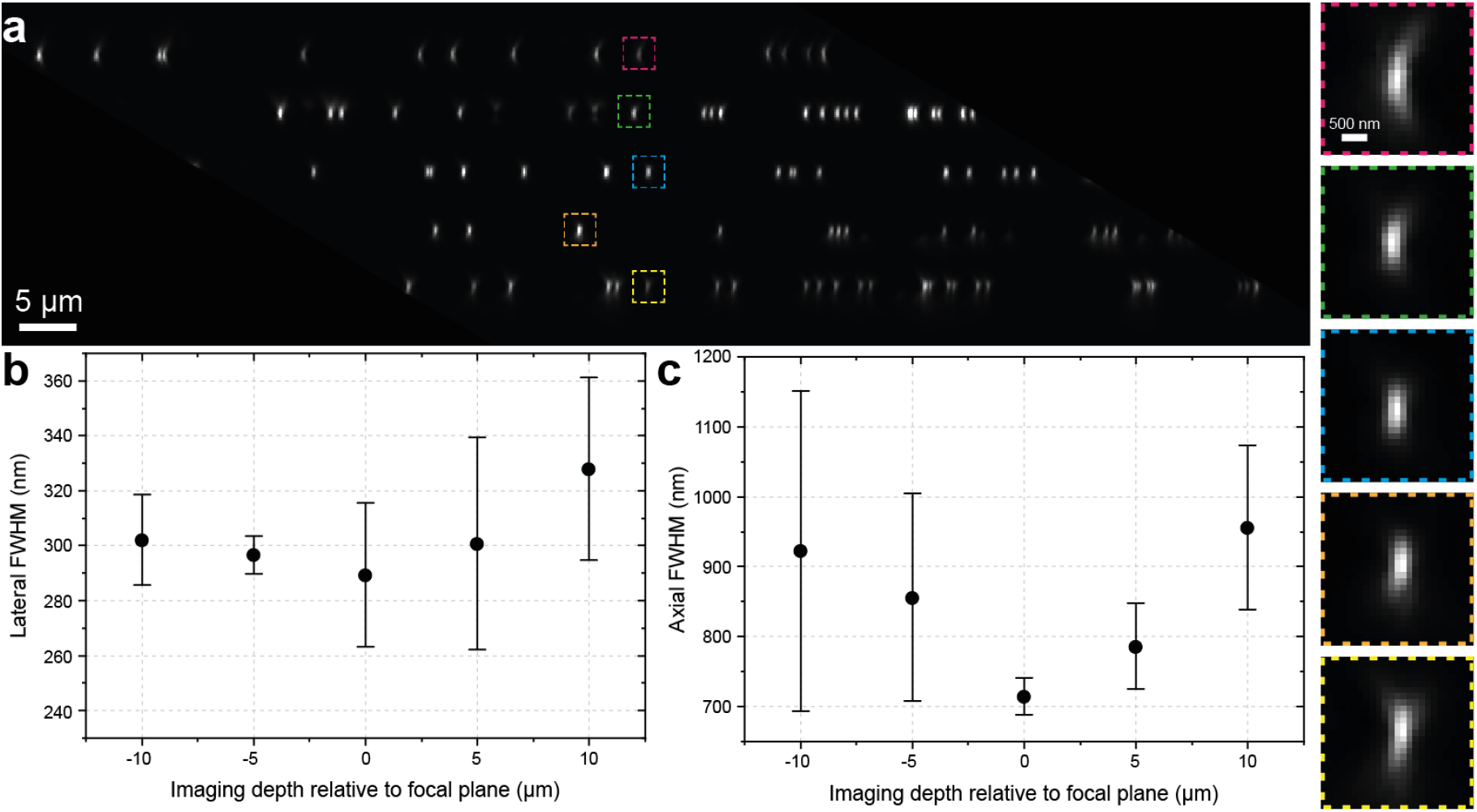
OPM PSF characterization. Volumes of 100 nm beads immobilized on cover glass were collected at depths between -10 to +10 μm with a 5 μm step, separately, and combined into a composite image with an XZ cross section shown in **(a)**. Boxed regions in (a) are shown to the right at larger scale. The mean lateral FWHM is shown in **(b)** while the mean axial FWHM is shown in **(c)** as a function of imaging depth relative to the focal plane.

### 3.4 Scanning unit characterization

We adopted a laser scanning approach for volumetric imaging as this method allows for high frame rates and does not mechanically agitate the sample. Most OPM implementations to date, rely on a scanning unit comprised of two scan lenses to maintain optimal remote focusing performance and a galvo mirror for beam steering [6,11]. In this scanning geometry, it is important that the galvo mirror is conjugated to the back focal plane of the primary objective. This restricts the motion of the objective close to an optimal position determined by the relay optics between the galvo mirror and the primary objective. Deviations from this optimal position result in a field-depended tilt of the excitation beam. This problem does not significantly affect other microscope modalities, such as confocal microscopy, that work with a diffraction limited excitation spot, but for OPM systems, where the light sheet can extend tens of microns above and below the focal plane, even small amounts of tilt can produce volumes with non-uniform illumination and poor signal to noise ratio.

To circumvent the above limitations and simplify microscope alignment, we opted for a two-mirror scanning geometry, where two galvo mirrors are positioned close to the primary image plane of the microscope base and are synchronously actuated to pivot the excitation beam around a point along the optical path after the tube lens [13,23]. The axial position of the pivot point can be controlled by appropriately adjusting the amount of angular displacement of each mirror as shown in Fig. 6a. Hence, by adjusting the ratio of the voltage signals that control the mirrors it is possible to compensate for the focusing motion of the primary objective during routine microscope operation and position the pivot point of the excitation beam at its back focal plane. This achieves tilt-invariant scans of the light sheet along the FOV regardless of the final focus position of the primary objective. Furthermore, this two-mirror scanning geometry produces a simpler setup with smaller footprint and avoids additional losses to the fluorescence signal compared to the traditional scan-lens/mirror configuration [24].

**Figure 6.**
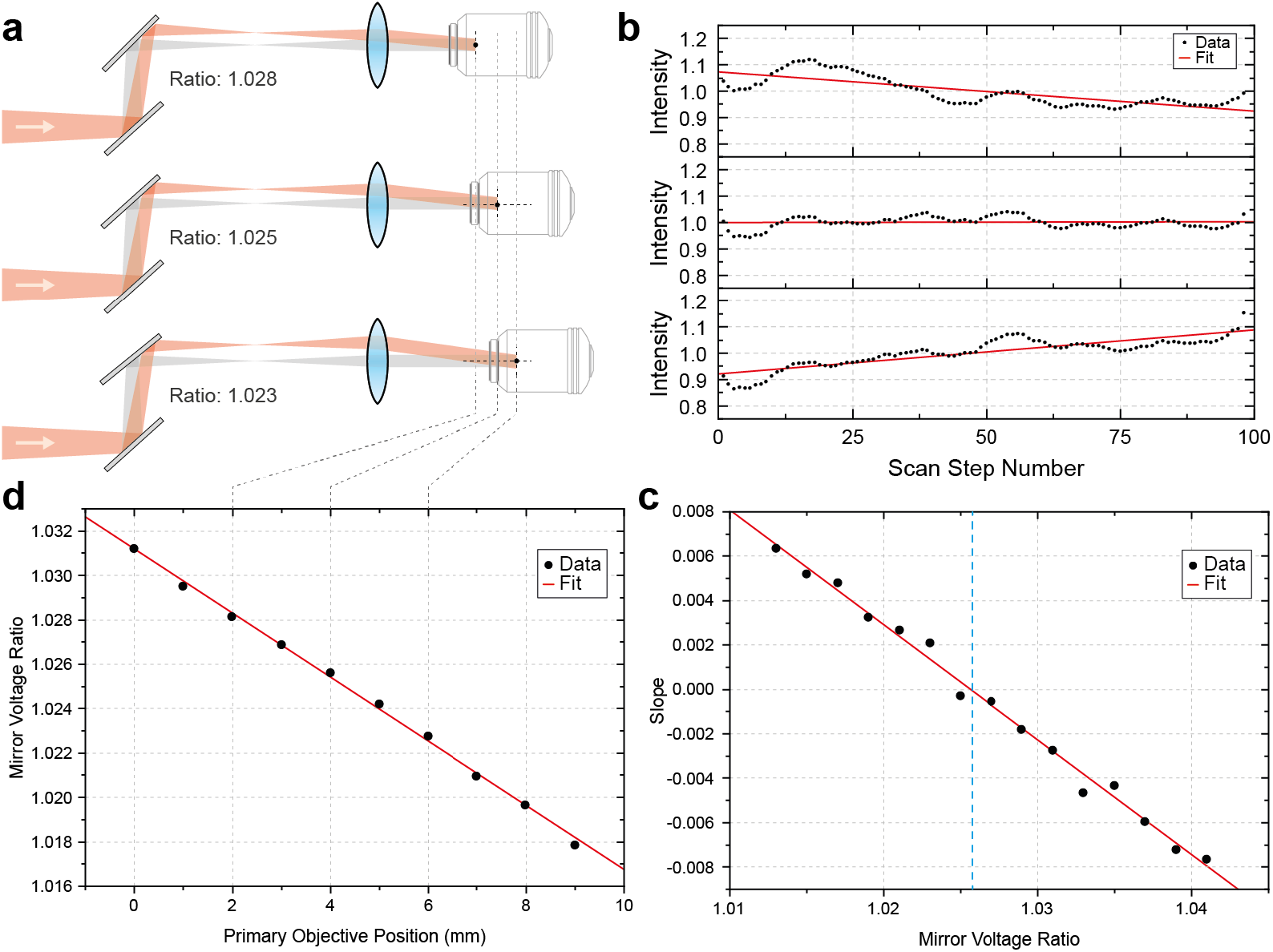
Scanning unit voltage ratio calibration. (**a**) The axial position of the pivot point of the excitation beam can be adjusted by changing the voltage ratio of the two scan mirrors to compensate for the motion of the primary objective during routine microscope operation. (**b**) Normalized intensity from a 100 nm bead embedded in a 2% agarose gel as a function of the steps taken by the two scan mirrors for three different voltage ratios between the waveform amplitudes driving the galvo mirrors for a given objective position. Each intensity vs. step data set was fit with a linear function to determine the slope and the ratio yielding the minimum slope was determined for that objective position (**c**). **(d)** The best ratio for each objective position was determined and fit with a linear function to generate a calibration for the mirror voltage ratio as a function of objective position.

To determine the optimal voltage ratio between the two mirrors for a given position of the primary objective we looked at scan-dependent variations in the fluorescence intensity collected from 100 nm beads uniformly distributed in an agarose gel for different mirror-voltage-ratios as described in Section 2.3 (**Fig. 6b**). We reasoned that when the mirrors are operated under the optimal mirror-voltage-ratio the light sheet is expected to maintain a constant tilt and provide uniform illumination of the oblique plane imaged by the camera at each step of the scan mirrors. In this scenario a linear fit of the summed intensity values as a function of mirror step would produce a slope that equals zero (**Fig. 6b**). Slope values above or below zero suggest non-uniform illumination of the imaged volume and consequently suboptimal mirror-voltage-ratios. In our setup this slope, which we use as a metric to evaluate the illumination uniformity of the scanned volume, appears to follow a linear relationship with the mirror-voltage-ratio. By performing a linear fit, we can therefore obtain the optimal mirror-voltage ratio for a given objective position as shown in **Fig. 6c**. By repeating this process for other objective positions, we can produce a calibration curve to map the entire travel range of the primary objective (**Fig. 6d**). It should be noted that adjusting the mirror voltage ratio to its optimal value as the primary objective is changing positions is straightforward in our setup and allows for seamless operation of the microscope. The objective Z position is read from the microscope base by the acquisition computer in real-time and the mirror-voltage-ratio is adjusted on the fly while the user adjusts the focal plane with the focus knob.

### 3.5 Microtubules dynamics

To demonstrate the capabilities of the microscope for high resolution fast volumetric imaging, we imaged microtubule dynamics in the follicular epithelium of Stage 7 *Drosophila* egg chambers. Microtubules are highly dynamic tubular structures with a characteristic diameter of 25 nm [25]. In *Drosophilla* follicle cells, microtubules are organized by apical microtubule organizing centres (MTOCS) and exhibit polarized growth extending from the apical to the basal side of the cell [26]. Previous work, either with whole microtubule labelling, or EB1 comets that track the growing end of microtubules, employed imaging rates in the range of ∼0.5-1 Hz [27,28]. Imaging at this speed is possible with other microscope modalities e.g., spinning disk, but is usually limited to a single plane. Therefore, given the 3D organization of the microtubule network, collecting data from a whole cell, instead of a single plane, would provide more useful datasets and a better understanding of the microtubule network dynamics [29].

Here, we imaged an oblique FOV of 100 × 20 μm which in cartesian coordinates corresponds to 100 × 10 μm (Y, Z) and is large enough to accommodate the entire dorsal-ventral axis of a Stage 7 *Drosophila* egg chamber up to a depth of ∼10 μm. This depth is sufficient to capture the entire volume of several epithelial cells, which, at this stage of development, have a height of ∼5 μm (Fig. 7a).

**Figure 7.**
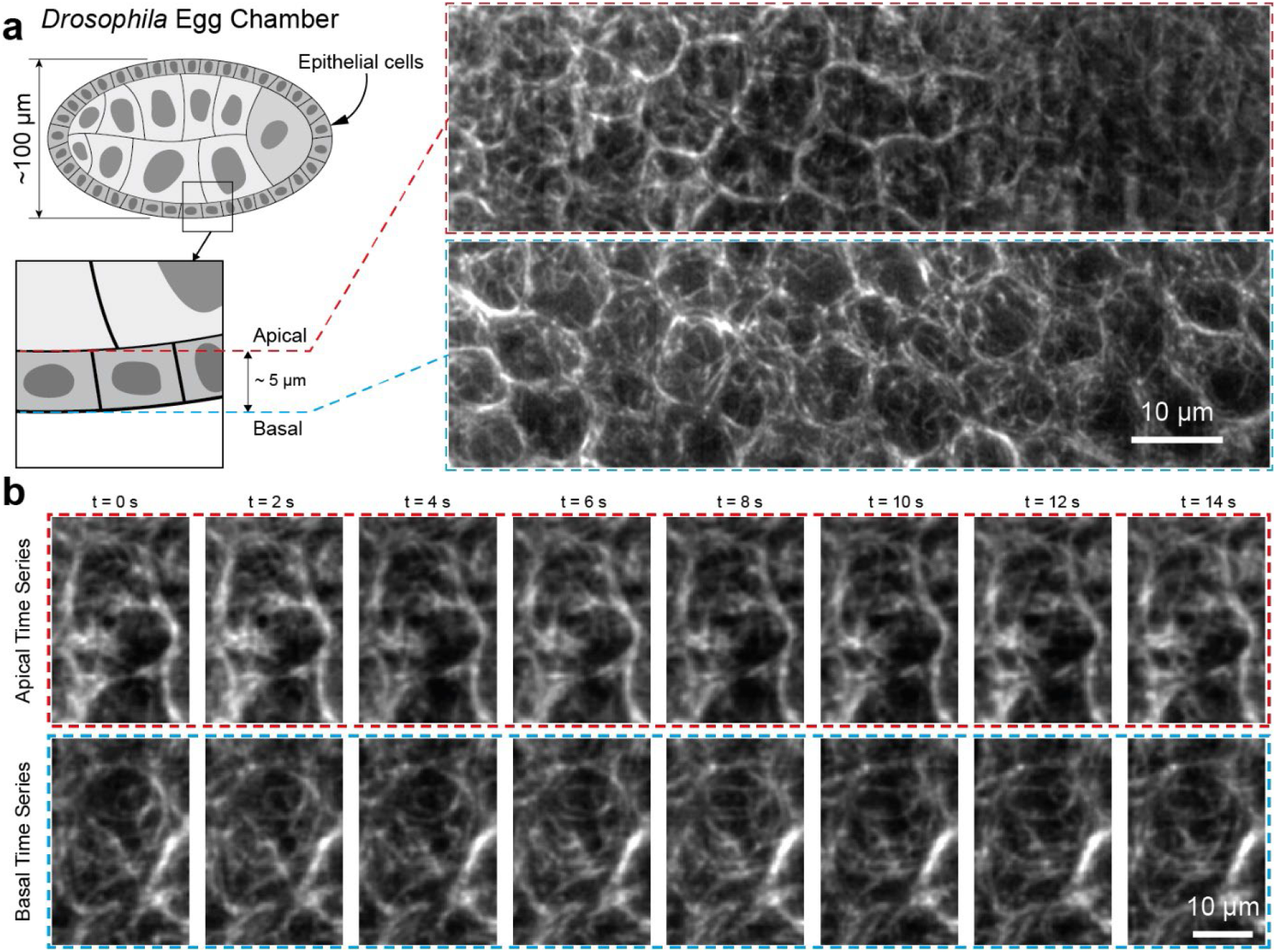
A volume in stage 7 Drosophila egg chamber with several epithelial cells was imaged at a rate of 0.5 Hz. (**a**) Overview images of the microtubule network, labelled with Jupiter-GFP, at the apical and basal planes of several epithelial cells. (**b**) Time series of a zoomed in region spanning a single epithelial cell showing the evolution of the microtubule network with the characteristic filamentous structures changing over time.

**Figure 8.**
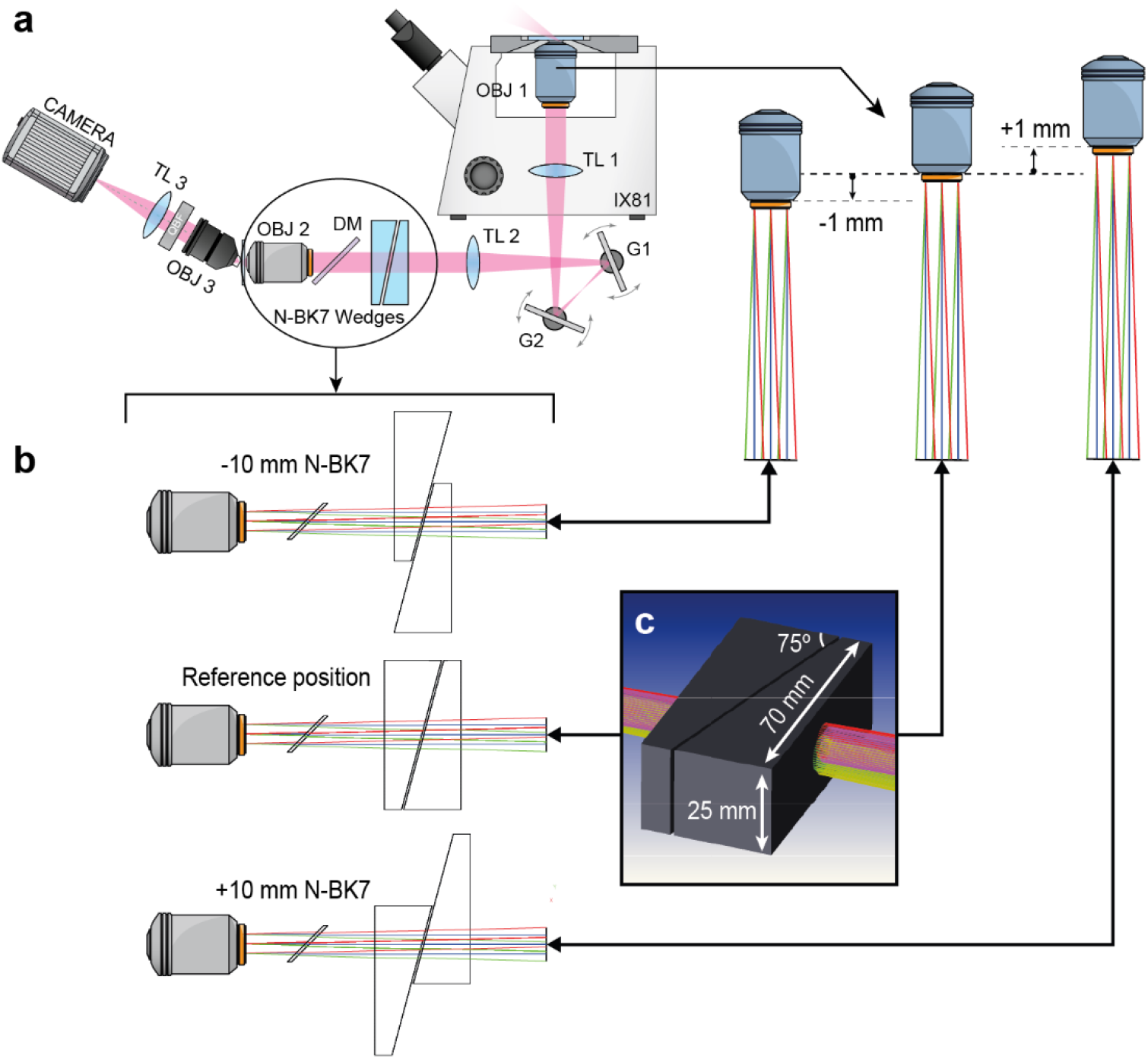
(**a**) System layout showing the position of the N-BK7 wedges in the optical path. (**b**) By moving the wedges, the amount of N-BK7 present in the optical path can be reduced or increased to compensate for displacements of the primary objective below or above its reference position during routine microscope operation. This maintains optimal conjugation between the primary and remote focusing objective during routine microscope operation and ensures uncompromised performance of the OPM microscope. It requires ∼10 mm of NBK-7 to compensate for 1 mm of primary objective displacement. The red, blue and green rays represent three different fields of a Zemax optical model from point sources at -85, 0, and +85 μm along the Y dimension of the field of view of the primary objective. (**c**) Shaded model of the N-BK7 wedges with relevant dimensions. OBJ: objective, TL: tube lens, G: galvo mirror, DM: Dichroic mirror

We were able to observe changes in the microtubule network with time and detect filamentous structures in both the apical and basal side of the follicle cells. The volumetric imaging speed mainly depends on the scan range, the step size and the sample brightness. Here, we used a camera exposure time of 10 ms and scanned the mirrors along the X dimension with a step size of 117 nm to achieve square pixels in the final deskewed volume and ensure Nyquist sampling. Under these conditions we imaged a total cartesian volume of 50 × 100 x 10 μm (X, Y, Z) at a rate of 0.5 Hz (Fig. 7b). However, by tuning the above experimental parameters, faster imaging rates can be achieved to match the speed requirements of different biological questions. For example, previous work tracking tips of microtubules suggested that an imaging rate of ∼1 Hz is required to observe rapid shrinking and growing events [25]. By halving the scan range from 50 to 25 μm, the imaging rate will double and volumes of 25 × 100 x 10 μm can be imaged at rates of 1Hz.

## 4. Conclusion

In this work we built and characterized an OPM microscope designed to work seamlessly with a commercially available microscope base. To support all the functionality offered by the microscope base, where the position of the objective lens is not fixed, we adopted a 2-mirror scanning geometry that offers a straightforward way to compensate for changes in the position of the objective lens during routine microscope operation. We showed that by adjusting the angular displacement of the mirrors in a calibrated way it is possible to capture 3D volumes with uniform illumination and high signal-to-noise ratio for every position of the objective lens within its travel range. This adjustability is not supported by the more traditional scan-lens/mirror scanning configurations. Furthermore, compared to the scan-lens/mirror, the proposed two-mirror scanning geometry provides improved light efficiency and a more compact footprint, offering potential advantages for all OPM designs, whether they employ a commercial base or not. We also showed that within the expected displacement range of the 100X, 1.35 NA objective lens from its design position, and for most practical applications, there is no significant effect on the resolving power, or the fidelity of the 3D data produced by the microscope to warrant further improvements in its optical path. To address applications beyond the scope of this work that require larger objective lens displacements or light sheets that illuminate volumes at greater depths, we proposed and discussed a modified optical path that incorporates a pair of glass wedges and allows for uncompromised performance of the microscope throughout the entire application space.

Given that microscope bases support standard sample mounting methods and various imaging modalities that enable non-expert users to perform a wide range of imaging experiments, we believe that this work describing the incorporation of an OPM modality in this familiar and user-friendly environment will further facilitate the adoption of this technology and make it available to a wider group of researchers.

### Appendix: Modified OPM optical layout with N-BK7 wedges

In section 3.2 we characterized the distortion of the remote focusing volume due to axial misalignment between the primary and the remote focusing objectives. We showed that for our application the expected distortions are below 0.5% and we did not take any additional steps to compensate for it. But, for other applications beyond the scope of this work we proposed a modified OPM layout that relies on a pair of N-BK7 wedges to compensate for the displacement of the primary objective during routine microscope operation. **Fig. 8** shows the details of this optical layout and illustrates the operation of the N-BK7 wedges.

## Funding

Wellcome Trust [080007, 203144, 203285, 207496]; Biotechnology and Biological Sciences Research Council [BB/P026486/1], and Cancer Research UK [A24823, C6946/A24843]. For the purpose of open access, the author has applied a Creative Commons Attribution (CC BY) public copyright license to any Author Accepted Manuscript version arising from this submission. Open access funding provided by the University of Cambridge.

## Disclosures

The authors declare no conflict of interest.

## Data availability

Data underlying the results presented in this paper are not publicly available at this time but may be obtained from the authors upon reasonable request.

## Notes

### Competing Interest Statement

The authors have declared no competing interest.

